# Sound-seeking before and after hearing loss in mice

**DOI:** 10.1101/2024.01.08.574475

**Authors:** Jessica Mai, Rowan Gargiullo, Megan Zheng, Valentina Esho, Osama E Hussein, Eliana Pollay, Cedric Bowe, Lucas M Williamson, Abigail F McElroy, William N Goolsby, Kaitlyn A Brooks, Chris C Rodgers

## Abstract

How we move our bodies affects how we perceive sound. For instance, we can explore an environment to seek out the source of a sound and we can use head movements to compensate for hearing loss. How we do this is not well understood because many auditory experiments are designed to limit head and body movements. To study the role of movement in hearing, we developed a behavioral task called sound-seeking that rewarded mice for tracking down an ongoing sound source. Over the course of learning, mice more efficiently navigated to the sound. We then asked how auditory behavior was affected by hearing loss induced by surgical removal of the malleus from the middle ear. An innate behavior, the auditory startle response, was abolished by bilateral hearing loss and unaffected by unilateral hearing loss. Similarly, performance on the sound-seeking task drastically declined after bilateral hearing loss and did not recover. In striking contrast, mice with unilateral hearing loss were only transiently impaired on sound-seeking; over a recovery period of about a week, they regained high levels of performance, increasingly reliant on a different spatial sampling strategy. Thus, even in the face of permanent unilateral damage to the peripheral auditory system, mice recover their ability to perform a naturalistic sound-seeking task. This paradigm provides an opportunity to examine how body movement enables better hearing and resilient adaptation to sensory deprivation.

## INTRODUCTION

Humans and other animals explore the world through body movement. For example, we process visual scenes by scanning our eyes over them (Yang, Lengyel and Wolpert, 2016) and we recognize physical objects by probing them with our hands and fingers (Lederman and Klatzky, 1987; Gamzu and Ahissar, 2001). In a similar vein, mice coordinate sniffing with locomotion to track down scents (Findley *et al*., 2021), vision with head movements to gauge distances (Johnson *et al*., 2021; Parker *et al*., 2022), and whisker movement with touch to recognize objects (Grant *et al*., 2009; Rodgers *et al*., 2021). Faced with the active nature of sensation, experimentalists may attempt to isolate sensory processing from motor control by physically restraining the subject (*e.g.*, with head fixation) or by using experimental designs that force a temporal separation in sensation, deliberation, and motor response (O’Connor *et al*., 2010). While separating sensation and action in this way has indeed illuminated fundamental principles of decision-making (Gold and Shadlen, 2007), our ability to extrapolate from restrained to free behavior is ineluctably limited. Moreover, the ubiquity of neural signals about body movement in sensory brain areas in head-fixed animals (Parker *et al*., 2020) suggests that physical restraint may not effectively isolate sensory processing and movement signals after all.

Although active sensing has primarily been studied in vision, touch, echolocation, and electrosensation, hearing is also an active sensorimotor process (Aytekin, Moss and Simon, 2008). In species with limited ability to move the ears independently, moving the head can optimize the collection of auditory information (Wallach, 1940; Perrett and Noble, 1997; Wightman and Kistler, 1999; Brimijoin, Boyd and Akeroyd, 2013; Yost, Torben Pastore and Dorman, 2020; Van Bentum, Van Opstal and Van Wanrooij, 2021; Higgins *et al*., 2023). For instance, we might turn to face a person in order to better hear them. Head movements can be used to favor the better ear in noisy environments like schoolrooms (Grange *et al*., 2018) or to adapt to hearing loss (Pastore *et al*., 2018, 2020; Gessa *et al*., 2022).

Neuroscientists often treat body movement as a confound whose effects must be cancelled out of sensory input by dedicated circuitry throughout the neuraxis (Crapse and Sommer, 2008; Requarth and Sawtell, 2011; Yu *et al*., 2016; Schneider, Sundararajan and Mooney, 2018). However, body movement can also create an opportunity to *enhance* sensory processing through neural circuits and computations that are less well understood (Chen, Murphey and MacIver, 2020; Mannella *et al*., 2021). These circuits and computations are difficult to study using only the tools available for human subjects; using mice as a model system enables a greater range of tools, such as cell type-specific monitoring and manipulation. Mice have excellent hearing, which they integrate with movement in order to survive and thrive. For instance, parent mice seek out their lost pups by the sound of their cries and bring them back to the nest (Ehret, 1987; Marlin *et al*., 2015; Dunlap *et al*., 2020).

In this project, we set out to understand how mice actively seek out sound and how they adapt to hearing loss. We developed a behavioral paradigm that we call sound-seeking, in which mice explore a multi-chambered arena to track down an ongoing sound source. Next, we surgically induced either unilateral or bilateral hearing loss in order to determine whether and how mice could compensate for disruption of peripheral input. After bilateral hearing loss, mice performed poorly and resorted to exhaustive search. In contrast, mice with unilateral hearing loss gradually regained good performance and relied increasingly more on an active motor sampling strategy. These results lay the groundwork for understanding how sensory and motor neural circuits enable successful navigation of complex auditory environments and recovery from sensory impairment.

## RESULTS

### The sound-seeking task

In order to understand how mice coordinate hearing and body movement, we began by developing an active auditory task that we call sound-seeking (Figure 1). In this task, mice navigate a multi-chambered arena to track down a sound source. Mice had restricted access to water in order to motivate them to earn water rewards for performing this task (Reinagel, 2018; Urai *et al*., 2021). During daily behavioral sessions, they were placed in an octagonal arena (Fig 1A-C) with a speaker mounted on each of its eight sides (*cf.* Ehret and Dreyer, 1984; Kacelnik *et al*., 2006). A reward port (or “nosepoke”) mounted below each speaker detected “pokes” (intrusions of the mouse’s nose) and could also dispense water in response to pokes that we wanted to reward. Transparent plastic walls partitioned the arena into eight peripheral chambers, each with its own speaker and nosepoke, as well as one central chamber from which all others could be reached.

**Figure 1.**
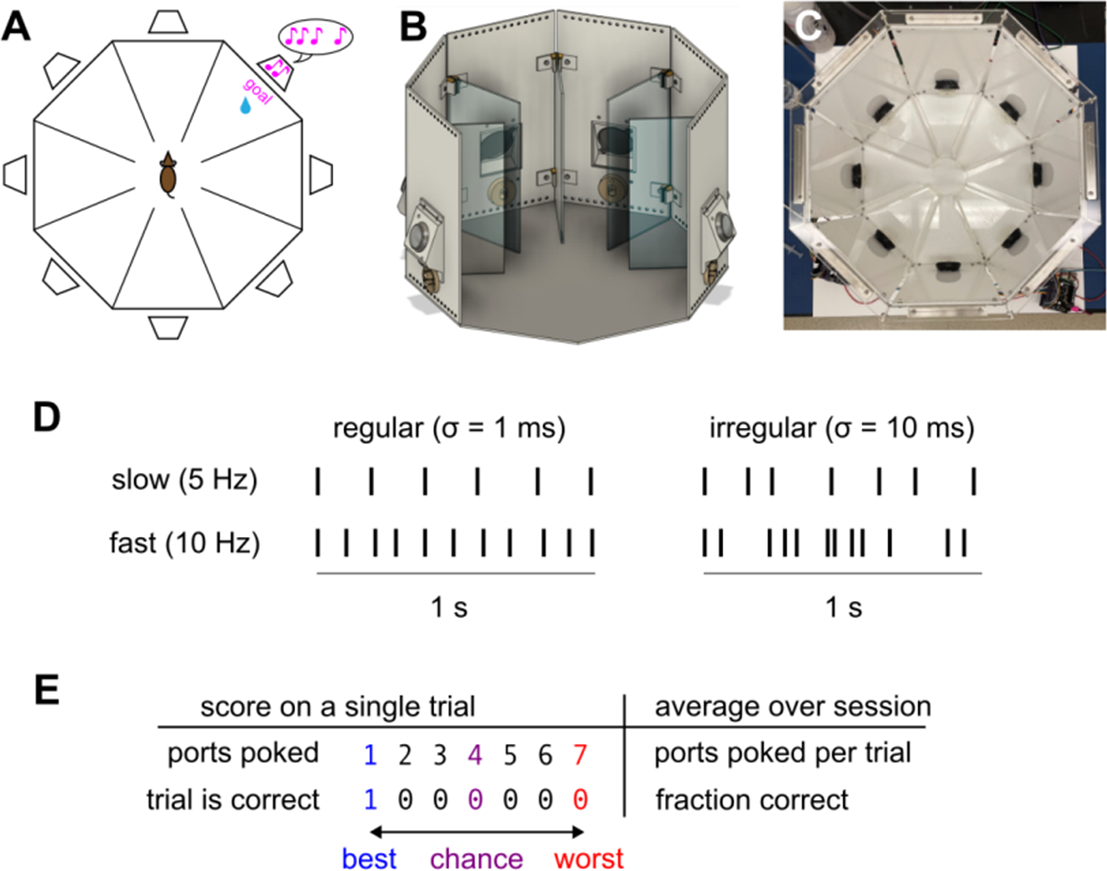
Freely moving mice learn to seek out a sound stream. **A.** Multi-chambered octagonal arena for sound-seeking. A speaker (trapezoid) was mounted on each wall above a nosepoke. On each trial, one speaker was randomly selected as the goal (pink). Mice received a water reward for poking the nosepoke below the goal speaker. **B.** CAD schematic of the arena with two external walls cut away to provide a view from the perspective of the mouse. Speakers are black; nosepokes off-white; and dividers transparent. **C.** Top-down photograph of the arena. Each side of the octagon was 15 cm and its longest diagonal was 40 cm. The entryway into each chamber was 5.5 cm wide. **D.** The auditory stimulus was an ongoing stream of intermittent noise bursts. The time between each noise burst was independently drawn from a gamma distribution with mean µ controlling the rate and standard deviation σ controlling the irregularity. **E.** A trial was scored as correct if the mouse poked the goal port before any other. We scored behavioral performance both as the fraction of correct trials and the mean number of ports poked per trial. Pokes into the previously rewarded port were never rewarded and did not affect scoring. Chance level performance (*e.g.*, random guessing) corresponds to 4 ports poked per trial and a fraction correct of 1/7 or 0.143. Perfect performance corresponds to 1 port poked per trial (the goal) and a fraction correct of 1.0.

Each session lasted approximately 25 minutes and comprised a series of trials. On each trial, one of the eight speakers (the “goal”) played a continuous stream of intermittent noise bursts. The auditory stimulus continued throughout the entire trial until the mouse poked the correct port (Fig 1D), in contrast to typical auditory localization tasks that use a single brief stimulus (Kavanagh and Kelly, 1987; Recanzone *et al*., 2000; Malhotra, Hall and Lomber, 2004; Kacelnik *et al*., 2006; Van Bentum, Van Opstal and Van Wanrooij, 2021; Town and Bizley, 2022). The mouse received a reward for poking the port below the goal speaker. Pokes into any other port triggered an error sound (a tritone with a base frequency of 8 kHz). Regardless of incorrect pokes, each trial lasted until the mouse poked the correct port and received reward. The mouse did not have to do anything in particular to initiate the next trial, which began 1 s after the previous reward. On each trial the goal speaker was chosen at random. However, the same speaker was never the goal on two consecutive trials because we did not want to reward the mice for perseverating in poking the previously rewarded port.

We quantified how well mice performed this task with two related metrics (Fig 1E). (Some mice perseverated in poking the previously rewarded port for several seconds after reward; thus, in calculating both metrics we disregarded pokes into the previously rewarded port, which could never be the goal.) First, we calculated the fraction of correct trials, defined as those on which the mouse poked the correct port before any other port.

Second, we calculated the mean number of ports poked per trial, which varied from 1 on a correct trial to 7 when the mouse poked every other port before the goal. (This metric could never reach 8 because we ignored pokes into the previously rewarded port, even if mice returned to it later in the trial.) To quantify performance we mainly relied on the number of ports poked, because this metric differentiates incorrect trials by how many incorrect ports were poked whereas the fraction correct metric treats all incorrect trials the same.

We trained five cohorts of mice under slightly different conditions (Table 1), not including a pilot cohort of mice used to test the hardware and determine initial task parameters (data not shown). As we optimized the task design, we sometimes changed between cohorts the mouse strain, water regulation protocol used for motivation, acoustic parameters, and arena geometry (Methods), until we arrived at a final choice of parameters in Cohorts 4 and 5. Despite this variation in parameters, learning was remarkably robust and consistent across animals. In particular, every mouse (n = 69) eventually learned the task well (see next section).

**Table 1.**
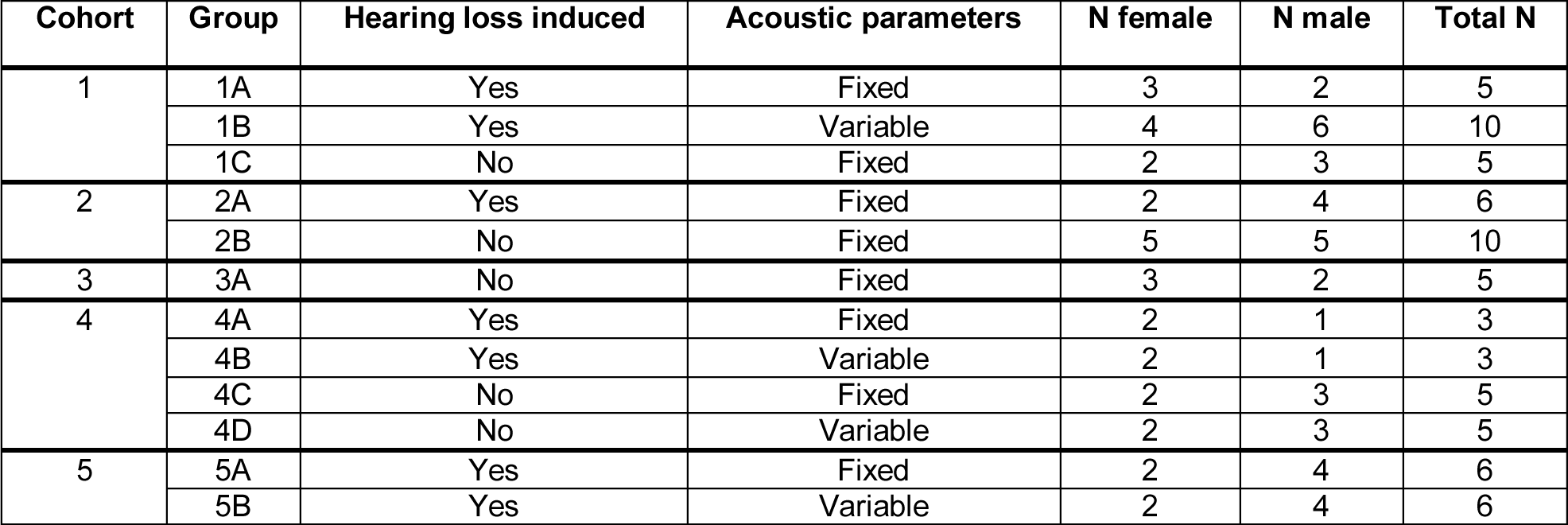
Mice used in this study. We trained five cohorts of mice for this study. Cohorts were divided into groups by whether the mice would receive hearing loss surgery or not, and whether the acoustic parameters would be varied or held fixed. Columns indicate number of female, male, and total mice. Cohorts 1-3 were used to refine the acoustic parameters and arena geometry; Cohorts 4-5 used the final values of acoustic parameters and arena geometry described in the text. Mouse strain and the water regulation used to motivate task performance also varied across cohorts as described in the Methods. Within each group, mice were always trained in the same way.

To determine which features of the sound affected mice’s decisions, some cohorts were divided into subgroups trained either with variable acoustic parameters (randomly chosen on every trial) or with fixed acoustic parameters (same on every trial). In all cases, the noise bursts were narrowband white noise with 10 ms duration and 3 kHz bandwidth. The intervals between noise bursts were drawn from a gamma distribution, which allowed us to independently vary repetition rate (mean number of noise bursts per second) and irregularity (standard deviation of the interval between noise bursts) (Fig 1D). For variable parameter mice in Cohorts 4 and 5, the center frequency was selected from the range 5 - 15 kHz, sound level from 65 - 90 dB SPL (measured at the center of the arena), repetition rate from 3 - 10 Hz, and irregularity from 1 - 100 ms. All of these acoustic parameters were constant within each trial but could vary across trials. For fixed parameter mice in those cohorts, every trial was the same: 4 Hz repetition rate, 31 ms irregularity, 70 dB SPL, 10 kHz center frequency. Cohorts 1-3 were trained with slightly different parameters (Methods).

### Despite varying conditions across cohorts, mice learned sound-seeking robustly and reliably

All mice (n = 69) began at chance but eventually learned the task (Fig 2A). Performance usually plateaued near 0.5 fraction correct and 1.8 ports poked per trial (meaning that on average expert mice poked 0.8 incorrect ports before the correct one). We defined early-learning and late-learning epochs as before and after the first session on which each mouse performed better than 2.5 ports poked per trial (red dots in Fig 2A). Mice achieved this criterion in a median of 11 training sessions (inter-quartile range: 8 - 16).

**Figure 2.**
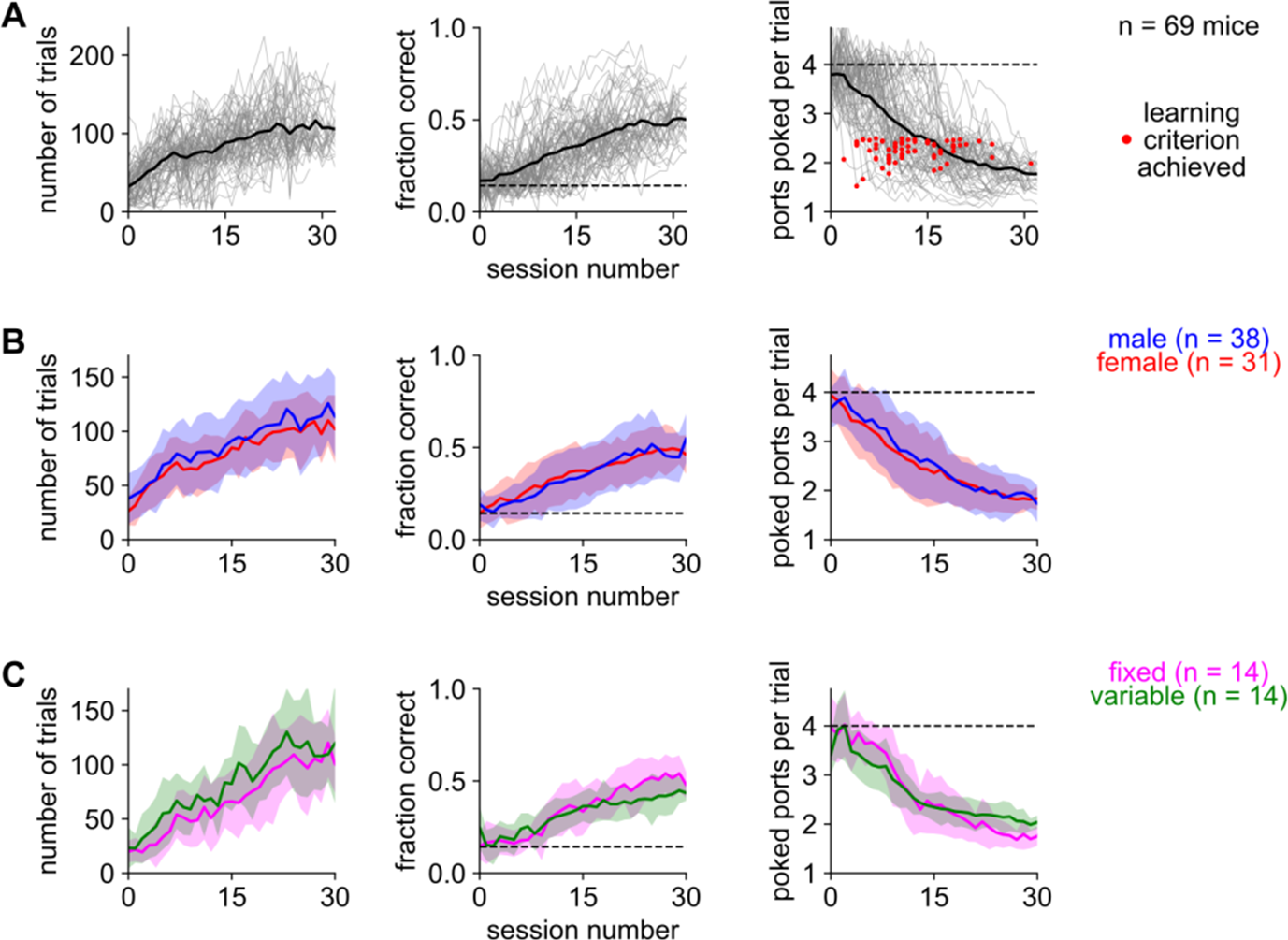
Male and female mice reliably learned either task variant. **A.** Performance of mice from all cohorts, quantified as number of trials (left), fraction correct (middle), and mean number of ports poked per trial (right) versus the number of training sessions. Gray lines: individual mice. Thick black line: average over mice. Horizontal dashed lines: chance performance. Red dots: first day each mouse achieved the criterion for good performance. **B.** Comparison of males (blue) and females (red). Males performed more trials (84.1 vs 76.6 trials; p < 0.01). Fraction correct and ports poked per trial did not differ (p > 0.05). **C.** Comparison of mice trained with fixed (magenta) or variable (green) acoustic parameters in cohorts 4 and 5. Mice trained on variable parameters did significantly more trials (p < 0.001) but performed slightly worse on the ports poked per trials metric (p < 0.01); there was no significant difference in the fraction of correct trials (p > 0.05). Statistical significance was assessed as the effect of group in a two-way (group + time) repeated-measures ANOVA. Shaded area: standard deviation.

Despite the variation in experimental parameters, including mouse strain and motivational paradigm (deprivation or unpalatability, see Methods), performance was roughly similar across Cohorts 1-5 (Supplemental Fig 1). Because of the multiple differences between cohorts, we cannot conclude whether the strain of the mouse or the motivational paradigm affected learning. However, each cohort included male and female mice that were trained simultaneously and identically, which enabled us to directly compare males and females (Fig 2B). Males performed 9.9% more trials than females, likely explained by their larger body size and increased need for water, but there was no significant difference in their fraction correct or ports poked per trial.

Also, we found only slight differences in subgroups trained on fixed or variable parameters (Fig 2C), suggesting that these two task variants are roughly equal in difficulty. All in all, mice of both sexes reliably learned either variant of this task in a few weeks of training.

### Mice were better able to seek out sounds with high repetition rate and low center frequency

Mice trained on variable acoustic parameters did similarly well across most of the parameter range that we tested (Fig 3). However, they did slightly better on trials with a high repetition rate, indicating they were able to take advantage of the higher information rate on these trials. They also did slightly better on louder than softer sounds, though in late learning performance appeared to plateau or even decline at high volumes (Fig 3B, third panel). Performance did not vary between irregular and completely predictable streams.

**Figure 3.**
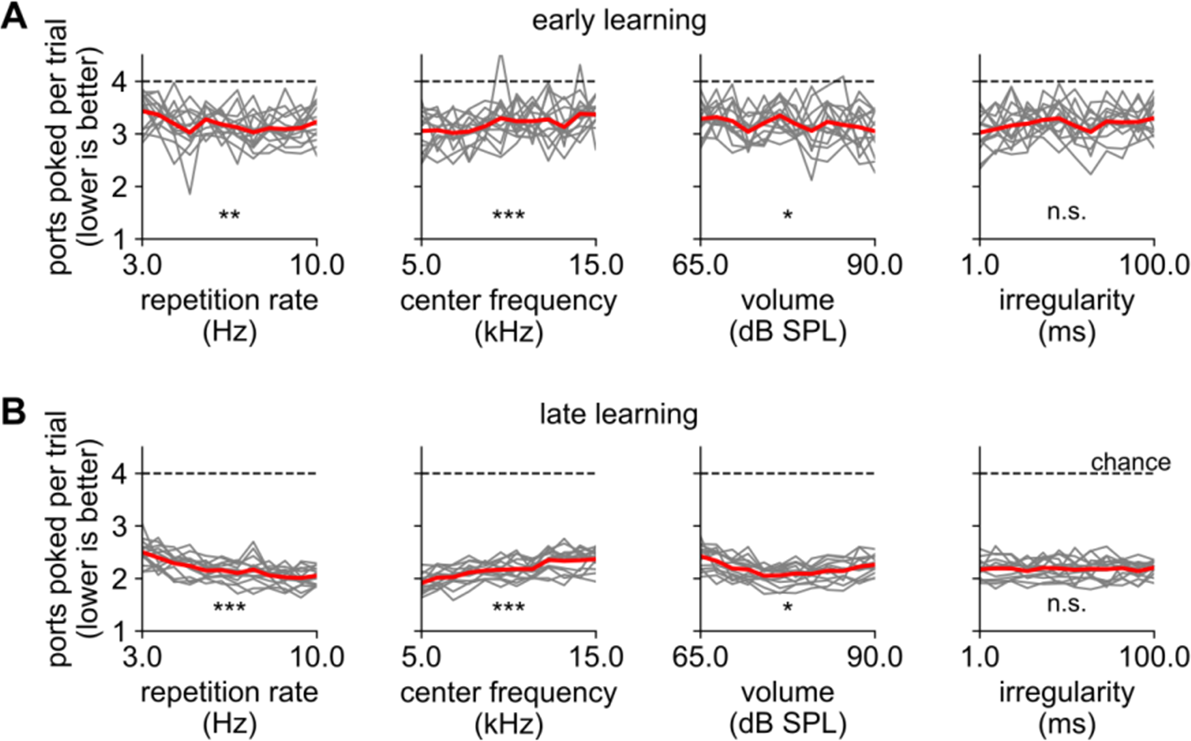
Mice seek out sounds roughly equally well regardless of their acoustic features. **A, B.** Performance as a function of acoustic parameters during early and late learning. Individual mice: gray lines; average: red; chance performance: dashed line. Mice performed significantly better for high repetition rates, low center frequency, and higher volume (p-value from Pearson’s correlation), but these effects were small. Irregularity had no significant effect on performance. All x-axes in these panels are logarithmic.

Surprisingly, mice were significantly better at seeking out low-frequency sounds. This was unexpected because mice localizing sound rely mostly on interaural level differences, which are more prominent at high frequencies (Lauer, Engel and Schrode, 2018). (The mouse head is too small to exploit the other main binaural cue, interaural timing differences.) In our task, perhaps low frequency sounds are more effectively restricted to single chambers, or they may reverberate differently. While reverberation is generally thought to impede localization, in some cases it can help (Hartmann, 1983). Alternatively or additionally, mice may rely on different cues for localizing sound while moving than they do in standard head-fixed conditions.

Notwithstanding these relatively small differences, mice performed roughly equally well for all sounds. In other words, within the range of parameters that we tested, task performance was mainly limited by errors that were unrelated to the acoustic properties of the sound.

### Resiliency to unilateral but not bilateral hearing loss

We next asked mice were affected by hearing loss and whether they could recover from it with sufficient practice. After mice had learned the sound-seeking task, we induced hearing loss in one or both ears by surgical extirpation (removal) of the malleus (Fig 4A-B). This surgery is a model of conductive hearing loss because the malleus and the rest of the ossicular chain conduct acoustic vibrations to the cochlea (Tucci, Cant and Durham, 1999; Xu, Kotak and Sanes, 2007; Lee *et al*., 2009; Salih *et al*., 2012; Mason, 2013); people often experience unilateral or bilateral conductive hearing loss as a result of ear infections, often in childhood (Moore, Hartley and Hogan, 2003; Whitton and Polley, 2011). Without the malleus, animals are not completely deaf but the intensity threshold for detecting sound is permanently elevated by 30 - 55 dB (Tucci, Cant and Durham, 1999; Xu, Kotak and Sanes, 2007). Mice were randomly selected for left, right, bilateral, or sham surgery, counterbalanced by sex. During bilateral surgery, the tympanic membrane was broken and the malleus removed on both sides. During sham surgery and during unilateral surgery on the spared side, the tympanic membrane and malleus were only visualized and not disturbed. In all cases, we used the same anesthesia, analgesia, and surgical scrub of the ear canal. We confirmed that unilateral and bilateral malleus removal affected hearing by measuring the acoustic brainstem response (ABR; Supplemental Fig 2) (Hall, 1992; Willott, 2006; Shaheen, Valero and Liberman, 2015).

**Figure 4.**
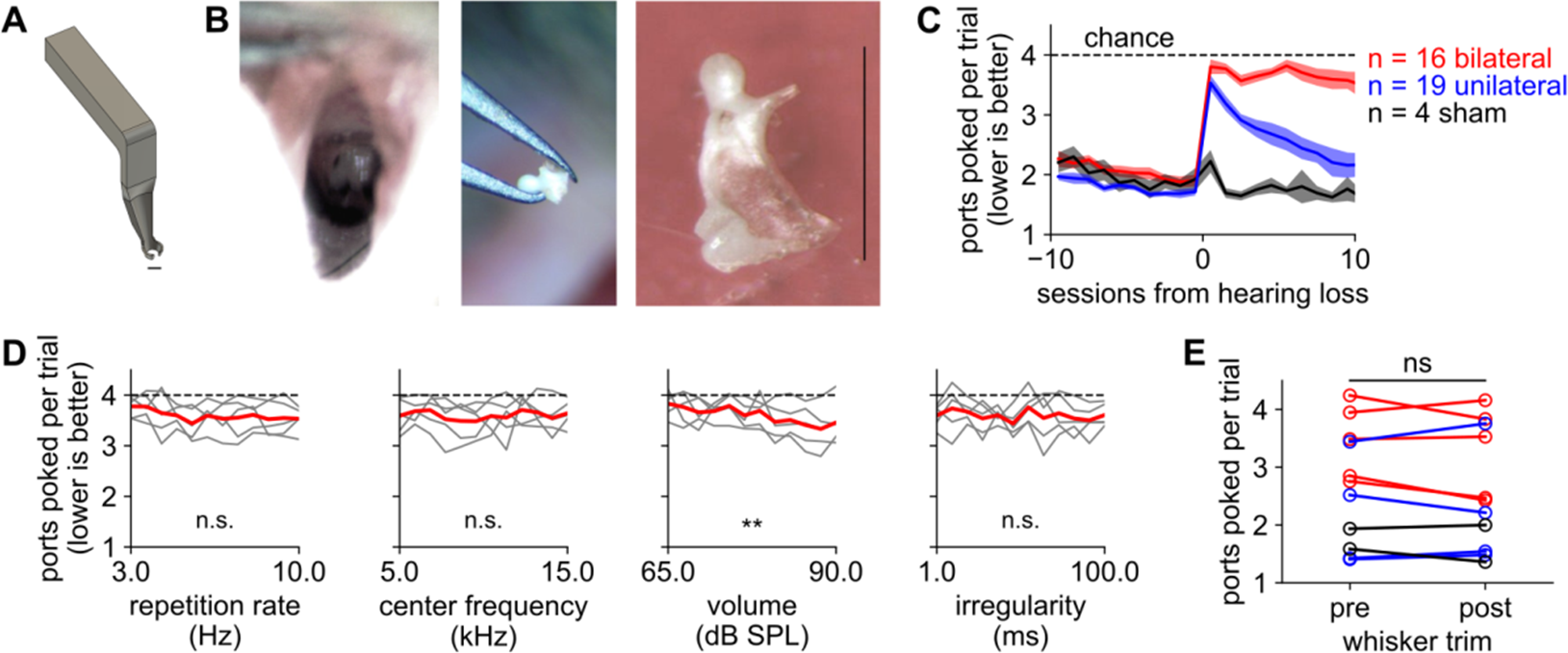
Sound-seeking performance recovers after unilateral but not bilateral hearing loss. **A.** Schematic of a 3d-printed miniature ear speculum for mice that we designed to visualize the middle ear. **B.** View into the ear canal (*left*), removal of malleus (*middle*), and high-magnification view of removed malleus (*right*). The manubrium has broken off this example malleus. Scale bars in right panel and in panel A are approximately 1.2 mm. **C.** Performance is significantly worse after hearing loss (p < 0.001 for bilateral and unilateral; p > 0.05 for sham) and significantly differs between groups after hearing loss (p < 0.001). Bilateral mice do not recover over time (p > 0.05) whereas unilateral mice do (p < 0.001). Dashed line: chance performance. We used ANOVA to assess statistical significance during recovery and between groups. To assess significance for an individual group, we used a paired t-test to compare before vs after hearing loss, and a p-value calculated from Pearson’s correlation with time during recovery. **D.** After bilateral hearing loss, mice performed significantly (p < 0.01) better on louder sounds than on quiet sounds, but only slightly. Conventions as in Fig 3. **E.** After whisker trim, performance does not change (n = 11 mice; p > 0.05; paired t-test). Colors as in panel B.

After hearing loss, we tested mice on the same sound-seeking behavior that they had previously learned. Mice with sham hearing loss showed either no change in performance or a minor impairment on the first session after surgery only (Fig 4C). Mice with bilateral hearing loss were profoundly impaired to near chance levels without improvement for up to 10 sessions after the surgery. Strikingly, mice with unilateral hearing loss were initially impaired but over 5-10 post-operative behavioral sessions their performance almost fully recovered (Fig 4C). Thus, mice rapidly regain their sound-seeking ability after losing sensory input on one side, but not if both sides are deprived.

Because malleus removal should raise the threshold of detection without completely deafening the mouse, we considered the possibility that bilateral mice would retain an ability to find loud sounds but not soft sounds. We tested this hypothesis in the subset of mice with bilateral hearing loss that were trained on the variable parameters version of the task, in which sound level and other auditory parameters were varied from trial to trial. These mice did show a small but significant increase in performance on trials with louder sounds (Fig 4D), raising the possibility that strong amplification (as in a hearing aid) might lead to further gains. However, the performance even on the loudest sounds (90 dB) was less than the performance on the softest sounds (65 dB) before hearing loss.

We also considered whether unilateral mice might have recovered their sound-seeking ability by learning to detect sound waves with their whiskers instead (Andermann *et al*., 2004; Yu, Graff and Hartmann, 2016). To test this, after at least 13 days had passed post-surgery, we cut off all of the macrovibrissae on a subset of mice. Their performance was unaffected by whisker trim, showing that they did not use whisker input to perform this task (Fig 4E). While it remains a possibility that mice could have learned some other strategy for sound-seeking, such as feeling floor vibrations, this would not explain why only the unilateral mice and not the bilateral mice relearned the task. The most likely explanation is that unilateral mice recovered by learning to rely on input from the spared ear (Kumpik and King, 2019), a strategy unavailable to bilateral mice.

### The acoustic startle response does not recover after hearing loss

We asked whether this recovery from unilateral hearing loss was specific to sound-seeking or would apply to all sensorimotor hearing tests, perhaps reflecting a generalized recovery of hearing ability over time. To do this, we assessed the auditory startle response, a brief whole-body muscular contraction in response to an unexpected sound at relatively high volume (Lauer, Behrens and Klump, 2017). This behavioral assay has been used to assess a variety of conditions; here, we simply use it to ask whether the mice can perceive and respond to sound, independent of the sound-seeking task. We measured auditory startle in Cohort 5 on three separate testing days over the same period of time during which we assessed their recovery on sound-seeking. Although auditory startle is usually measured with an accelerometer, we assessed it by tracking their body movements in video using the pose-tracking algorithm SLEAP (Pereira *et al*., 2022) so that in the future we can determine how these body movements are coordinated across the body (Fig 5A). This experiment was conducted in a dedicated behavioral chamber, not in the sound-seeking arena, with a background of continuous white noise at 60 dB SPL. The startle-eliciting stimulus was a 100 ms white noise burst with a level of 80 or 90 dB SPL. (We did not observe a difference between these two levels and so we combined them in this analysis.)

**Figure 5.**
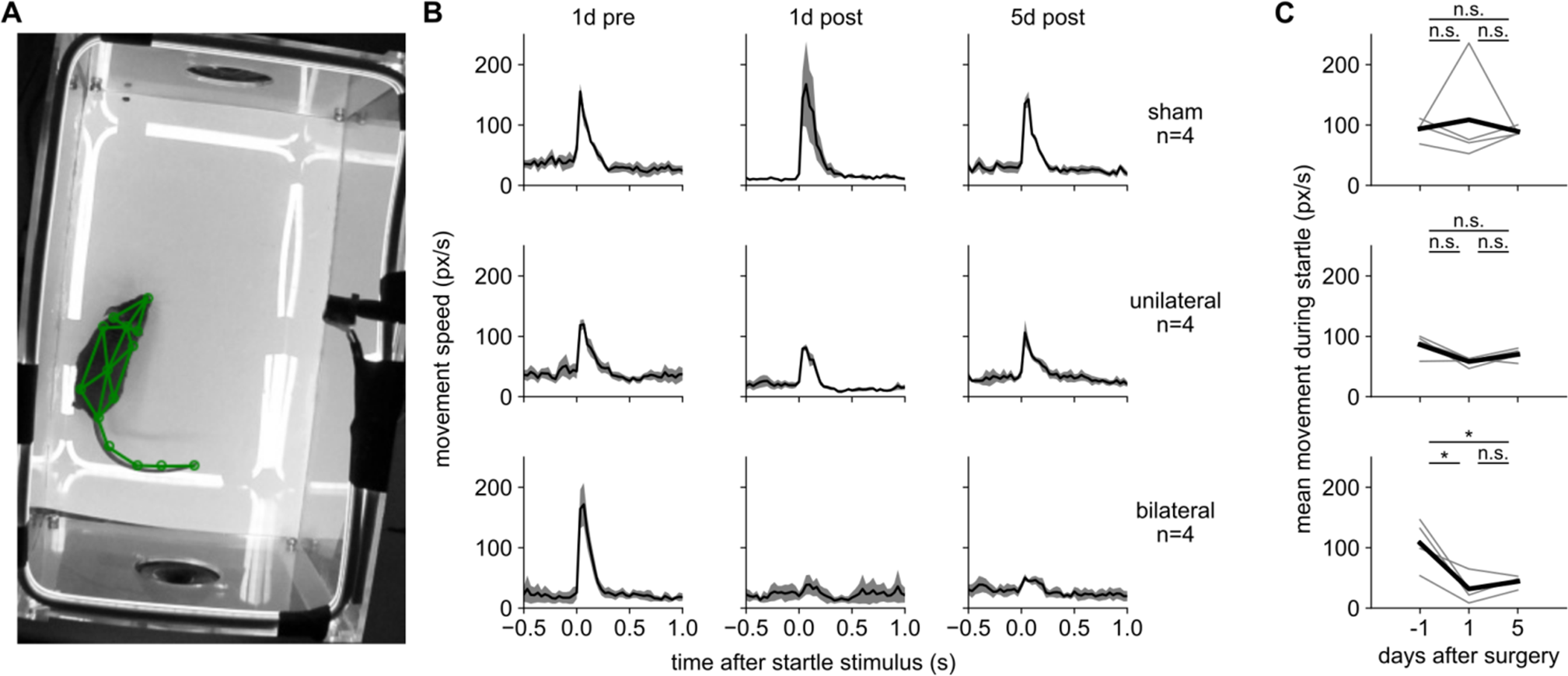
The auditory startle response is robust to unilateral hearing loss and does not recover after bilateral hearing loss. **A.** Example video frame showing rectangular arena with speakers at both ends, and mouse with body parts tracked in green. **B.** Acoustic startle response, quantified as full-body movement versus time from the onset of the startle-eliciting stimulus, at 1 day before, 1 day after, and 5 days after surgery (columns) for sham, unilateral, and bilateral mice (rows). Shaded area: SEM over individual mice. **C.** Acoustic startle response, quantified as the mean movement speed over the first five frames (167 ms) after the startle-eliciting stimulus. The response did not change significantly after sham or unilateral hearing loss, but it was significantly reduced after bilateral hearing loss (p < 0.05, paired t-test). Thin lines: individual mice. Thick line: mean over mice.

We measured the startle response in all groups on three days, one pre-operative to establish baseline and two post-operative to assess the effects of hearing loss (Fig 5B-C). After sham hearing loss, the startle response was generally unchanged at days 1 and 5 (although one outlier mouse was greatly enhanced at day 1 only). This result demonstrates that the startle response does not strongly habituate or diminish over time in our conditions. After bilateral hearing loss, the startle response was nearly abolished at post-operative days 1 and 5. After unilateral hearing loss, the startle response was nearly unaffected at days 1 and 5.

Because unilateral mice did not lose their startle response, we cannot assess their ability to recover it, but we can conclude that not all auditory behaviors are sensitive to unilateral hearing loss. While it is possible that additional testing of unilateral mice would have revealed a subtle impairment and/or recovery of the startle response, qualitatively this behavior was much less affected by unilateral hearing loss than sound-seeking was.

Because the bilateral mice did lose their startle response and did not recover it, we conclude that there is no generalized recovery of hearing over time after removal of the malleus (*e.g.*, through residual sensitivity of the cochlea accompanied by homeostatic regulation of gain in auditory afferents). Thus, the strong recovery after unilateral hearing loss that we observed in sound-seeking is unlikely to be explained by a generalized recovery of hearing ability. We speculate that the startle response may rely on brainstem circuits that require only input from a single ear, whereas recovery of sound-seeking may require time to recalibrate central auditory or motor circuits.

### Mice with unilateral hearing loss rely more on a motor strategy of checking each chamber

Because mice with unilateral hearing loss did not recover a typical acoustic startle response (indeed, full peripheral recovery from malleus removal is impossible), we next asked what motor strategies or behavioral changes might explain how they recovered their sound-seeking ability. For instance, mice might have taken a different path to the sound, taken longer to make their decision, or adapted their body posture. To quantify changes in motor behavior, we again used the pose-tracking software SLEAP to track the head and body position during sound-seeking of 18 mice (7 bilateral, 7 unilateral, and 4 sham from Cohorts 4 and 5) from initial task exposure through expert performance and recovery from hearing loss.

The general pattern is illustrated by representative trials in Fig 6A. Early in learning (first column), mice generally navigated the arena with a tortuous and self-crossing path, entering the same chambers and nosepokes multiple times. Later in learning (second column), mice were less likely to enter unrewarded nosepokes, less likely to enter the same chamber multiple times, and generally took a more direct path to the goal. After sham hearing loss, behavior remained stable (top row, right panels), but after bilateral hearing loss (middle row, right panels) mice made many more errors, often simply cycling in one direction around the arena and poking each port in turn. To quantify this pattern, we defined a “chamber entry” as the video frame on which the snout passed the internal dividers (Fig 6B). Duplicate entries (*i.e.*, re-entering a chamber that had already been visited on the same trial) significantly declined over learning and did not increase after hearing loss (Fig 6C). Cycling entries (*i.e.*, entering a chamber directly adjacent to the one just exited) significantly increased over learning and then sharply increased again after both bilateral and unilateral hearing loss but not after sham (Fig 6D). Thus, we find that naïve mice enter chambers unsystematically, often visiting the same chamber multiple times in a self-crossing trajectory, whereas expert mice cease to enter the same chamber more than once per trial, perhaps because they adopt a cyclical search strategy.

**Figure 6.**
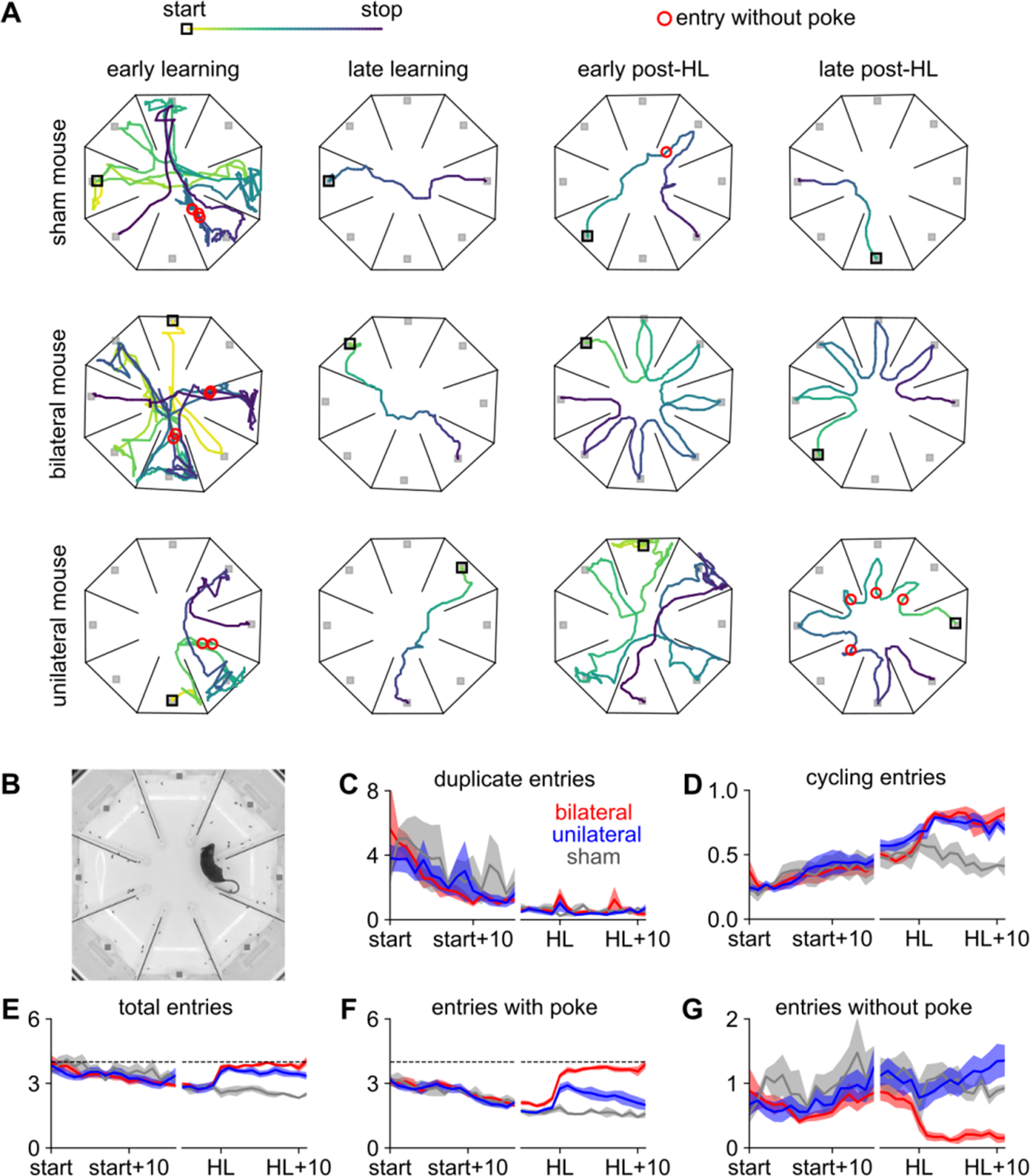
Over learning, duplicate and cycling entries decrease; after unilateral hearing loss, entries without poke increase. **A.** Example trials from different mice (rows) taken at four different timepoints over behavioral testing (columns). Colored line: mouse trajectory from early in trial (light colors) to late (dark colors), with the start marked as an open black square. Filled gray squares: port locations. Red circles: entries without poke (checks). **B.** Example frame showing the first entry without poke on the example trial in the bottom right of panel A. **C.** Mean number of duplicate chamber entries per trial significantly decreases over learning (p < 0.001). This does not significantly increase after hearing loss for any group (p > 0.05, t-test) or differ by type of hearing loss (p > 0.05). In panels C-G, the x-axis is broken into two epochs: a learning epoch comprising the first 15 sessions of training on the left, and a peri-hearing loss epoch comprising the 5 sessions before and 10 sessions after hearing loss surgery on the right. **D.** Mean number of cycling entries per trial significantly increases over learning (p < 0.001). These entries significantly increase after hearing loss (p < 0.01 unilateral; p < 0.001 bilateral; p > 0.05 sham) and are significantly more common in bilateral (p < 0.001) and unilateral (p < 0.001) mice than they are in sham mice. **E.** Mean number of entries per trial significantly decreases over learning (p < 0.001). These entries increase after hearing loss (p < .01 unilateral; p < 0.001 bilateral; p > 0.05 sham) to near the chance level of 4 (dashed line). Panels E-G exclude duplicate entries. Panels F and G further divide the chamber entries in panel E into two categories. **F.** Mean number of entries with poke per trial (essentially the same as plotting the number of pokes, Fig 4C). Entries with poke significantly decrease over learning (p < 0.001). Entries with poke increase after unilateral (p < 0.01) and bilateral (p < 0.001) but not sham (p > 0.05) hearing loss; they decrease over the recovery period for unilateral mice (p < 0.001) not bilateral mice (p > 0.05). **G.** Mean number of entries without poke per trial significantly increases over learning (p < 0.001). These entries sharply decrease after bilateral (p < 0.01) but not unilateral (p > 0.05) hearing loss. During recovery, entries without poke increase over time for unilateral mice (p < 0.001) and are more frequent than in either sham (p < 0.001) or bilateral (p < 0.001) mice. Throughout this figure, we used ANOVA to assess statistical significance during learning, during recovery, and between groups. To assess significance for an individual group, we used a paired t-test to compare before vs after hearing loss, and a p-value calculated from Pearson’s correlation with time during recovery.

Throughout learning, but especially later in learning and after unilateral hearing loss (bottom row, right panels), mice would briefly enter a chamber without poking the port (red circles in Fig 6A). We call this behavior “checking”, because we speculate that mice entered the chamber to check whether its speaker was playing sound. We hypothesize that checking could be a motor strategy to compensate for sensory loss. To quantify this observation, we counted the total number of chambers entered on each trial, discarded all duplicate entries and entries into the previous goal (which could never be the current goal), and divided the remaining entries into two categories: those that eventually culminated in a nosepoke (“entry with poke”) and those that did not (“entry without poke”, *i.e.* a check).

The total number of entries per trial began at around 4 (commensurate with entering chambers at random), decreased over learning, and increased to the chance level again after hearing loss (Fig 6E). After bilateral hearing loss, entries with poke increase while entries without poke (checks) almost entirely cease (Fig 6F-G, red line) – these mice enter the chambers in sequence and poke each port until they find the goal. In contrast, after unilateral hearing loss mice actually increase their entries without poke (checks; Fig 6G, blue line). That is, while both bilateral and unilateral mice visit unrewarded chambers at chance levels (Fig 6E), the unilateral mice increasingly often refrain from poking after entering an incorrect chamber. In other words, unilateral mice seem to have lost the ability to navigate directly to the chamber emitting sound, an ability which may rely on binaural cues; instead, they sample auditory space by physically moving through the arena, checking each chamber until they find the sound (Groh, 2014).

## DISCUSSION

In this study, we developed a sound-seeking task in which mice sought out a continuous sound source through ongoing movement of the head and body. Over training, mice learn to take a more efficient path to the goal that visited fewer chambers, often in a spatial sequence. After bilateral hearing loss, this spatial sequence became even more routinized as mice essentially visited each port in turn, leading to chance level performance. In contrast, mice with unilateral hearing loss were transiently impaired but recovered nearly full performance over time, although now increasingly reliant on a motor strategy of checking each chamber for sound without actually poking the port. We did not observe the same pattern of transient impairment followed by recovery in an innate behavior, the auditory startle response, suggesting that recovery on sound-seeking may rely on distinct neural circuits. We speculate that unilateral mice, no longer able to exploit binaural cues to navigate directly to the goal port, instead checked each chamber and relied on monaural input to release a nosepoke motor response. Overall, this work provides a platform to study how the brain can exploit motor or behavioral changes to build resilience to sensory impairment, with implications for rehabilitation and recovery in auditory and other modalities.

### Active sound-seeking versus passive sound localization

Sound-seeking challenges mice to integrate auditory input with movement of the head and body in order to find an ongoing sound source. Our results were presaged by early work in which mice were better able to localize ongoing sound if they could move toward it (Ehret and Dreyer, 1984). Sound-seeking sharply contrasts with what we call “passive sound localization” tasks, which are designed to separate body movement from hearing. For instance, passive studies might physically restrain the subject (Recanzone *et al*., 2000; Lauer, Slee and May, 2011; Coen *et al*., 2023), train them to hold still during the sound (Lomber and Malhotra, 2008; Lingner, Wiegrebe and Grothe, 2012; Town and Bizley, 2022), or use sounds so brief that they terminate before the subject can move (Kelly, 1980; Kavanagh and Kelly, 1987; Kacelnik *et al*., 2006; Nodal *et al*., 2010; Nodal, Bajo and King, 2012). For example, (Kelly, 1980) writes “Brief stimuli [were used] to eliminate the possibility of scanning movements of the head or body as a localization strategy.” Such approaches remove motor feedback in order to expose the precise sensory cues that animals can use to compute sound location (Goodman, Benichoux and Brette, 2013), as well as the underlying brainstem and cortical neural circuitry (Knudsen, 2002; Konishi, 2003; Grothe, Pecka and McAlpine, 2010; Keating and King, 2015; Lauer, Engel and Schrode, 2018; Middlebrooks, 2021). Our approach differs in three important ways.

First, rather than probe perception when movement is impossible, sound-seeking models what free animals might naturally do, such as during pup retrieval (Marlin *et al*., 2015; Dunlap *et al*., 2020). Fundamentally, accurate localization in any sensory modality requires information from at least two points in space. This information can be obtained with multiple sensors (*i.e.*, the two ears) or alternatively from a single sensor moving through space over time, but the latter is only possible if the subject is free to move and the stimulus is ongoing (Wallach, 1940). Rather than casting the process of decision-making as a feed-forward sequence of sensory processing, deliberation, and finally action selection, we favor a recurrent model in which perception and action bidirectionally guide each other (Dewey, 1896; Gibson, 1979; Ahissar and Assa, 2016; Yang, Wolpert and Lengyel, 2016; Chen, Murphey and MacIver, 2020). We noted that sound-seeking mice rarely stayed still but rather moved continuously; sensory input modulated body movement rather than driving it in lockstep.

Second, most studies of auditory localization take place in anechoic (echo-suppressing) chambers with minimal physical structure that could impede the passage of sound waves (Grothe, Pecka and McAlpine, 2010). Such a design is better for measuring the exact acoustic features that enter the auditory system, but ultimately it is not representative of the natural environment. In natural behavior, humans and animals live in reverberant, multi-chambered environments such as buildings and burrows (Hartmann, 1983; Hu and Hoekstra, 2017). Interpreting the resulting echoes is an important function of auditory neural circuitry (Lauer 2018). In our task, mice may exploit the physical structure of the arena by ducking briefly into each chamber in order to detect the sound level, as opposed to relying on binaural cues in the center of the arena. Exploiting the physical structure of the environment to ease neural computations is an adaptive strategy for cognition (Clark and Chalmers, 1998; Tytell, Holmes and Cohen, 2011).

Third, in our task mice registered their decision by physically approaching the sound, rather than by licking or pressing a lever. While the nosepoke itself is certainly not natural, one feature of natural behavior is approaching a sound (as in pup retrieval) or fleeing a sound (as in escape from predators). When perception and action are viewed as separate processes, the mode of response would seem irrelevant to sensory processing. However, from the perspective that the brain’s function is to produce adaptive action, the mode of response may drastically alter the underlying neural computations. For instance, classic work in bushbaby suggested that auditory cortex was required specifically to link auditory perception with an approach response (Ravizza and Diamond, 1974), as opposed to auditory localization per se (although see Kelly, 1980). Directing the eyes toward a sound enhances localization (Maddox *et al*., 2014), and it has even been suggested that auditory localization evolved to support directing the eyes toward important objects (Heffner and Heffner, 1992, 2016). During sound-seeking, the sensory problem of localizing sound is unified with the motor problem of orienting and moving toward it.

### Spatial strategies for foraging for rewards

Throughout learning, mice became more likely to cycle through the chambers in a clockwise or counterclockwise spatial sequence. Cycling in this way is reminiscent of traplining, a form of foraging in which animals revisit known potential sites for reward in a regular spatial order (Saleh and Chittka, 2007). According to the way we measured performance, cycling is a chance-level strategy, no more effective than randomly choosing ports. But from the animal’s perspective, cycling may be more efficient or cognitively easier because the same turning movement is employed over and over, and because the memory of the ports previously checked is embodied in the trajectory taken. In addition, the cycling strategy may be innately preferable for mice because it hugs the walls and avoids entering the open center of the arena (Rosenberg *et al*., 2021); thus, sound-seeking may require mice to balance phonotaxis with thigmotaxis.

Recent work demonstrated the gradual replacement of a spatial strategy with an auditory-guided strategy in a naturalistic behavior, likely guided by a projection from prefrontal to auditory cortex (Lu *et al*., 2023). While perseverative spatial biases limit performance in laboratory tasks with randomized reward locations, they are likely adaptive under natural conditions where rewards are rarely random and often reoccur. Even primary sensory cortex can encode choice and reward history over trials (Rodgers *et al*., 2021). While on the one hand parietal cortex encodes trial history and causally drives perseverative responses (Akrami *et al*., 2018), it is also possible that other brain regions encode trial history for the purpose of overcoming these biases and producing the uncorrelated responses necessary for success in laboratory tasks. The prefrontal cortex is one candidate structure, as it both encodes auditory responses and is necessary for task switching (Floresco, Block and Tse, 2008; Rich and Shapiro, 2009; Rodgers and DeWeese, 2014). Sound-seeking provides an experimental platform for understanding how neural circuits update, balance, and select distinct strategies throughout learning and in response to sensory loss.

### Recovery from hearing loss

Hearing loss lowers the signal-to-noise ratio (SNR) at the input to the auditory system. Adaptive filtering in neural circuits can extract the most signal out of this weakened input (Keating and King, 2013)). However, the data processing inequality means that no amount of neural processing can restore hearing in full; informally, “turning up the volume” can only do so much because it amplifies noise as well as signal (Chambers *et al*., 2016; Asokan *et al*., 2018; Balaram, Hackett and Polley, 2019). The same logic limits the theoretical efficacy of hearing aids. Moreover, the cognitive load associated with difficult auditory processing might worsen or accelerate dementia (Griffiths *et al*., 2020; Paciello *et al*., 2021).

However, body movement is a potent way to enhance auditory processing that could help the healthy and hearing-impaired alike. For instance, in very noisy environments one might turn one ear toward the speaker to better capture their voice; this response can be either innate or learned, and it can benefit both healthy and hearing-impaired listeners (Grange *et al*., 2018). To find out which smoke alarm is chirping, one is more likely to explore the house room by room than to sit stationary and compute interaural localization cues (Groh, 2014). Finally, interaural time and localization cues are identical for a sound exactly in front and exactly behind the listener, but this ambiguity can be instantly resolved by moving the head to a new angle, which will generate very different cues for the two previously ambiguous cases (Perrett and Noble, 1997; Wightman and Kistler, 1999; Gessa *et al*., 2022). These movement strategies improve SNR by moving the sensor rather than by implementing complex algorithms to process noisy data.

Movement is a natural, innate strategy for improving our hearing. Translationally, a better understanding of these movement strategies and how they can be taught through physical therapy could provide a less invasive, less expensive, and faster treatment plan. Scientifically, this leads to the counter-intuitive idea that recovering from sensory disorder may rely on plasticity in motor circuits. The sound-seeking task provides a platform to study how networks of sensory and motor brain areas work together to enable the ongoing interaction between perception and action that we call behavior.

## METHODS

All animal experiments were conducted under the guidance and approval of the Emory University Institutional Animal Care and Use Committee under protocol number 202100154.

### Behavioral training

Mice were purchased from Jackson Laboratories or bred in a pathogen-free barrier facility (additional details given in Supplemental Table 1). In either case they were kept in single-sex cages after weaning. We generally provided enrichment such as running wheels and manzanita sticks, and we usually avoided selecting mice with signs of barbering or who were resistant to handling. We always attempted to use males and females in equal proportion, though this was not always possible given the vagaries of mouse breeding.

Training began with 2-5 days of handling, in which the experimenter habituated the mouse to being picked up and weighed. Then we began water restriction, which could be of one of two types as specified in Table 1. For water restriction, the water bottle was removed from the cage, and mice only received water during their daily behavioral training. We monitored how much water they received during the experiment, and provided a calibrated amount of water as needed to ensure that they always received their daily water ration of at least 4% of their body weight per day. We used a system of weight thresholds and daily health checks to ensure that mice remained healthy on this regimen, and in some cases increased their daily ration as needed to maintain a healthy weight and disposition. For palatability training, mice had unlimited access to water containing 1.5% - 3% citric acid (Sigma) in the standard home cage bottle. Citric acid is a food preservative that lends water a sour and unpalatable flavor without affecting the subject’s health, even over long-term consumption (Reinagel, 2018; Urai *et al*., 2021). These mice were motivated to perform the same task for the same water rewards because the pure water tasted better than their home cage water. All mice began at 1.5% citric acid but were increased to higher concentrations if their trial count was substantially lower than typical. We typically lowered the water reward size as mice became better at the task to encourage them to perform more trials.

After 1-2 days on this water regimen, mice began “pre-training” which lasted about 3-7 days (one session per day) and taught them to use the nosepokes. The first stage of pre-training was “manual poke training” in which they were introduced to the behavioral arena. In this stage we placed ∼0.1 mL of water inside the nosepokes using a syringe so that they would learn to associate the nosepokes with reward. After mice learned to reliably obtain their daily ration from the nosepokes, they began “automated poke training”. In this stage, no sounds were ever played, and every poke into a nosepoke triggered the release of a small water reward (∼5 - 15 µL), with the exception that the same nosepoke never delivered a reward twice in a row. Mice quickly learned to visit nosepokes serially, often in a cycling pattern. Once they learned to reliably obtain their daily ration in this way (typically 2 - 5 days), pre-training ended and they began training on the sound-seeking task described in the main text. At this point in their training, the median mouse was 91 days old (IQR: 83 - 115).

The behavioral system was controlled with our fork (https://github.com/Rodgers-PAC-Lab/autopilot) of the Autopilot behavioral control system (Saunders and Wehr, 2019). Briefly, the experimenter used a graphical user interface on a desktop PC that communicated with a Raspberry Pi microcontroller over a wireless network. This “parent” Raspberry Pi chose the stimulus parameters on each trial and communicated them to four “child” Raspberry Pi. Each child Pi controlled two speakers (either Multicomp Pro MCPCT-G5100-4139 or Almencla Ultrasonic Tweeter 2-inch Waterproof Piezo Horn) using a HiFiBerry Amp2 audio card and two Autopilot nosepokes using custom-built printed circuit boards. The child Pi communicated to the parent Pi every nosepoke and reward, and the parent Pi reported these values to the graphical user interface on the desktop PC, where they were stored in an HDF5 file for later analysis.

We used an ultrasonic microphone to measure the sound power at every frequency, and we corrected the amplitude of the sound for the frequency response of the sound card and speaker to ensure that the sound pressure level was consistent across frequencies. Some earlier cohorts were trained without this calibration procedure. The arena was constructed of acrylic and custom 3d-printed parts. The behavioral training was conducted inside a custom-designed sound-attenuating chamber (Item24). Water rewards were delivered to the nosepoke by opening a solenoid valve (The Lee Co LHDA05311115H).

### Speaker calibration

We calibrated the end-to-end frequency response of our audio system using an industrial microphone (Hottinger Brüel & Kjær 2829 and 4939). The microphone was placed in the center of the arena pointing directly at the wall containing the speaker. During calibration we recorded full-bandwidth white noise, measured the spectrum of the received acoustic power, and calculated the correction factor needed at each frequency to reach a uniform spectrum. During the behavioral session, we applied this correction factor to the sounds in software before sending them through the audio card. After this procedure, the frequency response of our audio system was within +/-10 dB over the range 5 kHz to 80 kHz, and within +/-5 dB over the range used for behavioral testing (5 - 20 kHz).

The sound level of the click stimulus used for the auditory brainstem response (ABR) was measured using a different microphone (BAFX 3370) that applied A-weighting rather than a flat frequency response. Therefore we report this sound level in units of dBA instead of dB SPL.

### Hearing loss surgery

To induce hearing loss, we surgically removed the malleus on one or both sides of the head. Briefly, animals were administered pre-operative extended-release buprenorphine and either carprofen or extended-release meloxicam. They were anesthetized with isoflurane throughout the procedure. We used aseptic technique and ensured surgical depth of anesthesia with the toe pinch reflex. The hair around the ear was shaved or removed with depilatory cream. The skin of the ear canal was disinfected with three alternating washes of Betadine and 70% ethanol. Under a surgical microscope, we visualized the tympanic membrane using fine forceps and/or a custom-designed 3d-printed miniature ear speculum. The tympanic membrane was broken and the malleus removed with fine forceps, but the other ossicles were left in place. Finally the ear canal was flushed with sterile saline. For sham surgeries, the procedure was identical up to and including visualization of the ossicles and flushing with saline, but the tympanic membrane and all ossicles were left intact and in place. The experimenters training the mice were blinded to the form of hearing loss experienced by each mouse.

### Acoustic Brainstem Response (ABR) measurement

We measured the physiological function of the inner ear and auditory nerve by measuring the ABR (Hall, 1992). Mice were anesthetized with isoflurane and placed belly-down with the head centered in a low-profile silicone anesthesia mask. A speaker was placed 12 cm to the left or right of the mouse facing directly toward the head. The same speaker and HiFiBerry sound card was used as in the behavioral training with the same calibration procedure, but the stimulus was a 0.1 ms click at either 60 or 70 dBA. The ABR measurement was repeated at least twice with the speaker on the left side and again with the speaker on the right side. We measured the ABR on four separate occasions: immediately before hearing loss surgery, immediately after hearing loss surgery, and following the 6^th^ and 7^th^ post-operative behavioral sessions. The results were quite variable even within the same mouse, and so for each mouse we chose the post-operative measurement with the strongest and cleanest ABR, as assessed by an experimenter who was blinded to the type of hearing loss.

We measured the ABR using an integrated circuit for multi-channel differential recording of biosignals (Texas Instruments ADS1299) and custom-build hardware to report the response to a computer. Regardless of the position of the speaker, the positive (*i.e.*, non-inverting) and negative (inverting) electrodes were 30-gauge stainless steel needles placed in the skin just ventral to or just posterior to the intertragal notch. The ground electrode was the same type of needle placed in the skin above the neck or at the base of the tail. This system is designed to collect full-bandwidth data at a sampling rate of 16 kHz. We also measured the speaker voltage on another channel. We noted some cross-talk with the speaker voltage on the other channel of the amplifier and removed this by finding the best linear filter to predict neural voltage from speaker voltage and then removing this prediction from the neural signal. We post-processed the data by filtering it above 100 Hz, removing outlier trials (*i.e.*, those in the top 5% of voltage excursion or standard deviation), and averaging approximately 750 repetitions.

The first peak in the ABR (“Wave 1”) originates from the auditory nerve (Hall, 1992). Because the positive (non-inverting) electrode was on the left ear, Wave 1 is expected to be negative for sounds from the left and positive for sounds from the right. For presentation purposes we inverted the response to sounds from the left so that Wave 1 would be expected to be positive in all cases.

### Startle test

We assessed basic auditory-motor function before and after hearing loss using the acoustic startle reflex (Lauer, Behrens and Klump, 2017). The same mice that were trained on sound-seeking were placed in a different behavioral chamber, a 17 by 18 by 27.5 cm rectangular prism constructed entirely of clear acrylic plastic and with a speaker mounted on both ends (Fig 5A). Sounds were generated by the same type of Raspberry Pi, HiFiBerry sound card, and speaker as used in the sound-seeking task. The startle-eliciting sound was a 100 ms burst of broadband white noise at either 80 or 90 dB SPL. These two levels were interleaved, but we observed that the resulting behavior was fairly consistent so we averaged them indiscriminately. Continuous 60 dB white noise played throughout the experiment. The interval between sounds was a random time interval between 5 and 30 seconds. The entire procedure was repeated on five days: the third from last and the last behavioral session before surgery, and the first, fifth, and seventh sessions after surgery. The data presented here comprise only the session before and the first and fifth sessions after surgery.

Before, during, and after sound presentation we took video of the mouse in the startle chamber at 640×480 resolution and 30 frames per second (White Matter e3vision). We tracked the position of the mouse’s head nose, limbs, trunk, and tail using the SLEAP algorithm. In this way we measured the “startle” response of the mouse after each sound, which is a full-body coordinated muscular contraction. Specifically, we measured the distance moved by each body part in pixels between each adjacent frame, averaged across all body parts, and aligned this motion to the onset of the sound.

### Data analysis

All data analysis was conducted in Python using the packages IPython (Perez and Granger, 2007), pandas (McKinney, 2010), numpy (Van Der Walt, Colbert and Varoquaux, 2011), scipy (Virtanen *et al*., 2020), statsmodels (Seabold and Perktold, 2010), and matplotlib (Hunter, 2007). We used SLEAP (Pereira *et al*., 2022) to measure body position in the startle test. The data and code necessary to regenerate the figures shown here will be shared upon publication of this manuscript.

In Fig 6, we assessed statistical significance during learning as the effect of time in a one-way ANOVA; before and after hearing loss as the effect of surgery in a two-way ANOVA (pre/post x group); during recovery as the effect of time in a two-way ANOVA (day x group); and between groups during recovery as the effect of group in a two-way ANOVA (day x group) that only included the groups in question.

Throughout the manuscript, * indicates p < 0.05, ** p < 0.01, and *** p < 0.001.

## ACKNOWLEDGMENTS

We would like to acknowledge Dan Polley for guidance on the choice of hearing loss procedure, Anita Devineni and Robert Liu for comments on the manuscript, and the following funding sources: NIH/NIDCD R21 DC019711, a Whitehall Foundation Research Grant, a NARSAD from the Brain and Behavior Research Foundation, Emory University undergraduate research fellowships (OH and JM), NIH/NINDS T32 NS096050 (LW and AM), and the Federal Work-Study program.

## AUTHOR CONTRIBUTIONS

KAB, MZ, and CR developed the hearing loss surgical procedure. WNG designed the hardware and low-level data acquisition software for collecting ABR measurements, and CR and RG performed the ABR measurements and analyzed that data. JM and MZ performed hearing loss surgeries. VE, JM, RG, OH, EP, and MZ trained the mice on the sound-seeking task. JM performed the startle test experiments and trained the SLEAP model for those videos. RG constructed the behavioral equipment and designed the ear speculum. CB, LW, and CR trained the SLEAP model for sound-seeking. AM reviewed the literature and helped to edit the manuscript. CR developed the initial version of the behavioral task, supervised the project, wrote the manuscript, and generated the final version of the analyses.

## SUPPLEMENTAL FIGURES

**Supplemental Table 1.**
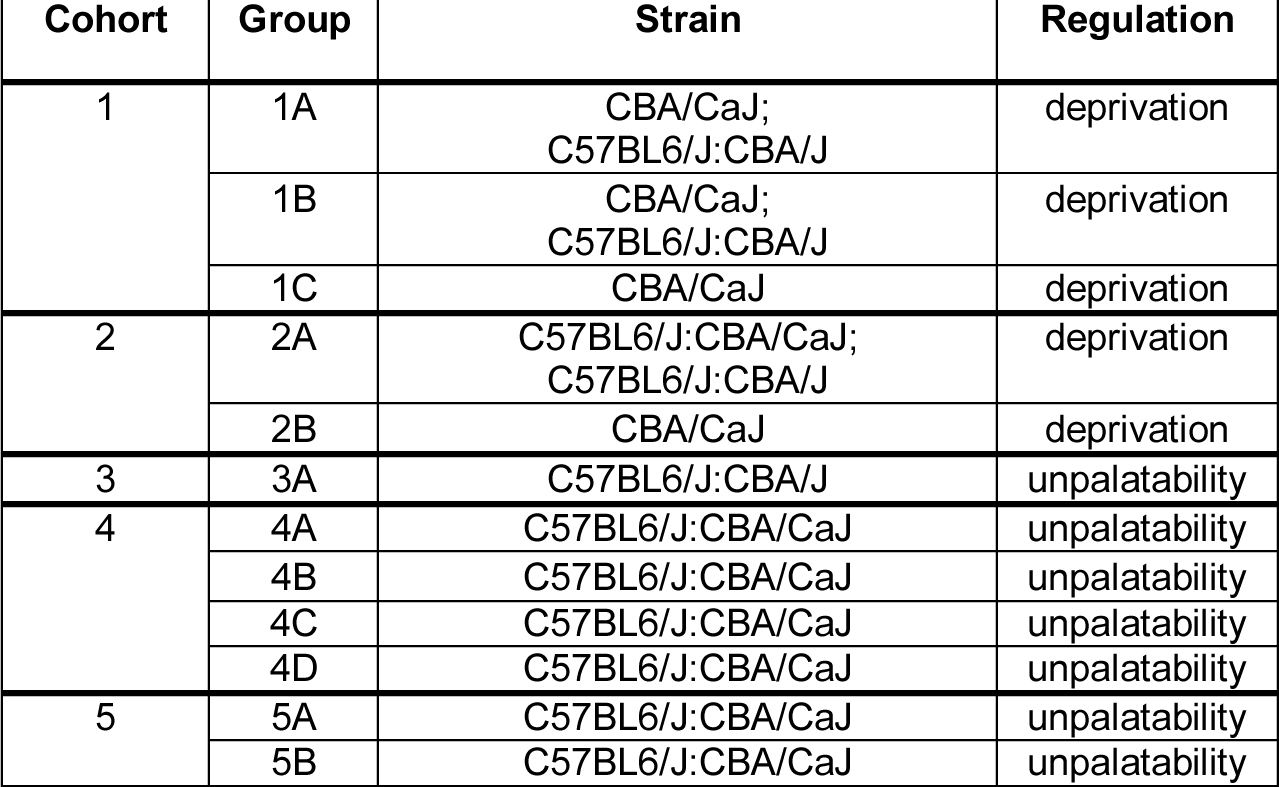
Mice used in this study.

This table provides additional information about the mice used in this study, with cohort number and group names matching Table 1 in the main text. The column “Strain” indicates the strain of mouse used in each cohort, with a colon indicating an F1 hybrid of two strains. Groups 1A, 1B, and 2A comprised mice of different strains or hybrids, which are separated with a semicolon and a linebreak in the table. Mice were either purchased directly from Jackson Laboratories or bred in our animal facility. All breeder mice were purchased from Jackson Laboratories, with stock numbers 000664 (C57BL6/J), 000654 (CBA/CaJ), and 000656 (CBA/J).

The column “Regulation” indicates the water regulation paradigm used. “Deprivation” means that no water was available in the home cage. “Unpalatability” means that mice had ad libitum access in the home cage to water containing 1.5-3% citric acid, which lends a slightly sour flavor. In all cases, rewards during the behavioral task were fresh water from a water purification system (Millipore). Mice were monitored daily for health while on any form of water regulation.

The arena geometry was varied slightly throughout Cohorts 1, 2, and 3. For instance we tried different lengths and heights of the dividers between chambers, and we switched from mounting the speakers on the inside of the wall to a recessed holder instead. Additionally we tried longer inter-trial intervals up to 5 s, switched from one type of speaker (Multicomp Pro MCPCT-G5100-4139) to another (Almencla Ultrasonic Tweeter 2-inch Waterproof Piezo Horn), and refined our speaker calibration procedures. Cohorts 4 and 5 were trained with a consistent arena geometry, finalized speaker calibration, and an inter-trial interval of 1s.

**Supplemental Figure 1.**
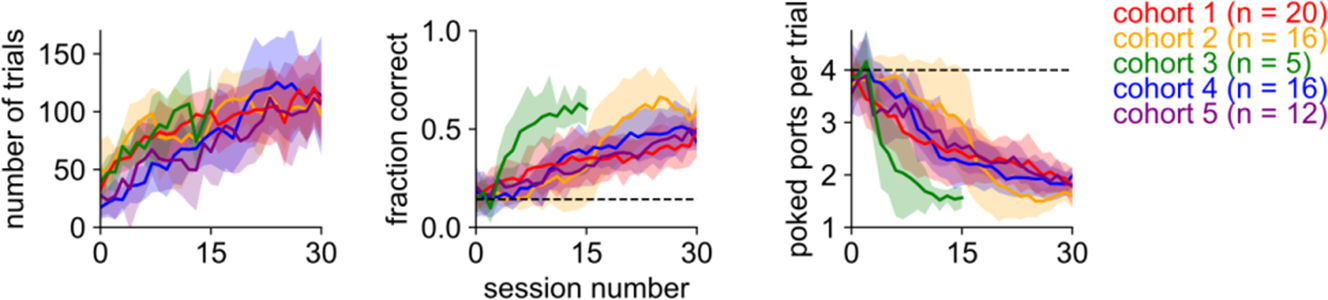
Learning across sex and cohort. Learning was variable between cohorts of mice but all cohorts reached a roughly similar level of final performance.

**Supplemental Figure 2.**
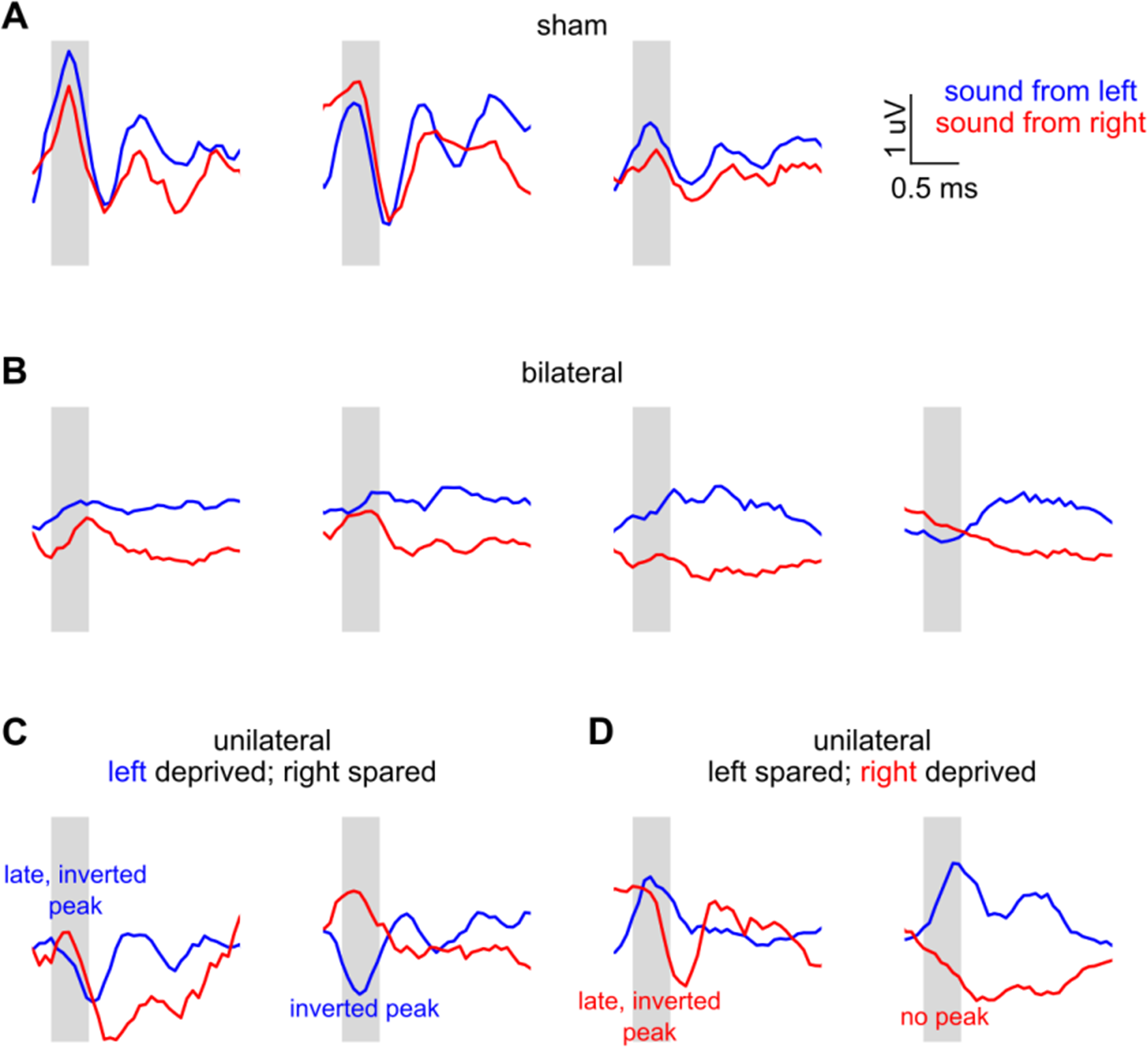
Acoustic brainstem response (ABR) after hearing loss. **A.** ABR following sham hearing loss surgery (control) in response to a click from left (blue) or right (red). This is a differential voltage measured at the left and right ear, averaged over about 1000 clicks. Gray shaded box indicates 2.0 - 2.3 ms after the click, the expected time of the first peak (“Wave 1”). The ABR signal has been inverted for sounds from the left to make Wave 1 positive. All three mice show strong, positive peaks within the shaded box for sounds from both sides. **B.** ABR following bilateral hearing loss. Wave 1 is attenuated, delayed, or abolished. **C.** ABR following unilateral (left) hearing loss. Wave 1 is delayed or inverted (negative) for sounds from the left (blue). Because of the differential recording procedure, the inversion likely represents a positive response at the other (spared) ear. **D.** ABR following unilateral (right) hearing loss. Wave 1 is delayed or inverted (negative) for sounds from the right (red).

